# High-frequency hearing is required for generating a topographic map of auditory space in the mouse superior colliculus

**DOI:** 10.1101/2021.10.21.465327

**Authors:** Yufei Si, Shinya Ito, Alan M. Litke, David A. Feldheim

## Abstract

A topographic map of auditory space is a feature found in the superior colliculus (SC) of many species, including CBA/CaJ mice. In this genetic background, high-frequency monaural spectral cues and interaural level differences are used to generate spatial receptive fields (RFs) that form a topographic map along the azimuth. Unfortunately, C57BL/6 mice, a strain widely used for transgenic manipulation, display age-related hearing loss (AHL) due to an inbred mutation in the Cadherin 23 gene (*Cdh23)* that affects hair cell mechanotransduction. To overcome this problem, researchers have used young C57BL/6 mice in their studies, as they have been shown to have normal hearing thresholds. However, important details of the auditory response characteristics of the SC such as spectral responses and spatial localization, have not been characterized in young C57BL/6 mice.

Here we show that 2-4-month C57BL/6 mice lack neurons with frontal auditory RFs and therefore lack a topographic representation of auditory space in the SC. Analysis of the spectrotemporal receptive fields (STRFs) of the SC auditory neurons shows that C57BL/6 mouse SC neurons lack the ability to detect the high-frequency (>40kHz) spectral cues that are needed to compute frontal RFs. We also show that crossing C57BL/6 mice with CBA/CaJ mice or introducing one copy of the wild-type *Cdh23* to C57BL/6 mice rescues the high-frequency hearing deficit and improves the topographic map of auditory space. Taken together, these results demonstrate the importance of high-frequency hearing in computing a topographic map of auditory space.

**Significance Statement:** Despite the strain’s age-dependent hearing loss, C57BL/6 mice are widely used in auditory studies because of the development of transgenic reporter and Cre lines in this genetic background. Here we examined the topographic map of auditory space and spectrotemporal properties of neurons in the SC of C57BL/6 mice. We found an early-onset high-frequency hearing deficit that results in the loss of SC neurons with frontal RFs and, consequently, an absence of a topographic map of auditory space. These findings stress the importance of high-frequency hearing in generating spatially restricted receptive fields and serve as a caution to researchers that doing auditory-related research using the C57BL/6 genetic background may not be representative of true wild-type mice.

## Introduction

The ability to discern the location of a sound is critical for survival. The superior colliculus (SC) is a midbrain structure that is important for sound localization and is the only brain area that contains a topographic map of auditory space. This map is aligned with the maps of visual and somatosensory space. SC circuitry uses this information to determine saliency and initiate motor commands for orientation and escape behaviors (Shang et al., 2015; Sparks et al., 1990; Stein & Clamann, 1981; Wang et al., 2015; Wei et al., 2015).

We recently performed an extensive physiological analysis of auditory responsive neurons in the SC of awake behaving mice (Ito et al., 2020). We found that auditory neurons in the mouse SC have spatially restricted RFs that form a topographic map of azimuthal space, that interaural level differences (ILDs) and the spectral filtering of the sound by the head and pinnae (spectral cues) are important for generating spatial RFs, and that there is a gradient in the relative weight of these features across the anterior-posterior (A-P) axis of the SC. Anterior neurons (that have frontal RFs) depend more on contralateral high frequency (40-80 kHz) spectral cues, while posterior SC neurons (that have lateral RFs) rely predominantly on ILDs.

Our previous study used CBA/CaJ mice as a model, which have been shown to have good hearing (Zheng et al., 1999) and is used in many auditory studies (Linden et al., 2003; Portfors et al., 2009; Portfors & Felix, 2005; Willott et al., 1988). Numerous auditory studies have been conducted using C57BL/6 mice to take advantage of the genetic tools made available in this background (Mao et al., 2006; Zhang et al., 2005). However, this strain has age-related hearing loss (AHL) (Henry & Chole, 1980; Willott et al., 1988; Zheng et al., 1999). This is due to an inbred mutation in the cadherin 23 gene (*Cdh23*), which affects cochlear hair-cell tip link formation and stability (Noben-Trauth et al., 2003). Studies have characterized the hearing threshold of C57BL/6 mice and found that hearing starts to decline in mice older than 3 months (Park et al., 2010; Zheng et al., 1999). In light of this, auditory researchers typically use C57BL/6 mice before or during early adulthood, assuming AHL has not yet taken effect (Kuchibhotla et al., 2017; Trujillo et al., 2013; Zingg et al., 2017). However, whether a young C57BL/6 mouse has the equivalent hearing ability as a CBA/CaJ mouse, and whether it contains a topographic map of auditory space in the SC, remain unknown.

To address these questions, we determined the spatial RFs of individual auditory neurons in the SC and found that 2-4-month C57BL/6 mice lack auditory responsive neurons with frontal RFs; therefore, C57BL/6 mice do not have a topographic representation of auditory space in the SC. We then determined the spectral temporal receptive fields (STRFs) of auditory neurons and found the SC of C57BL/6 mice lacks a high-frequency (40-80 kHz) response, consistent with the hypothesis that the high-frequency spectral cues are required for neurons to form frontal RFs in the SC. In an effort to rescue this high-frequency deficit, we measured the spatial RFs and STRFs in the first generation offspring from a cross of CBA/CaJ and C57BL/6 mice (F1(CBA/CaJ × C57BL/6)), as well as the C57BL/6 mice introduced with one copy of the wild-type *Cdh23* (B6.cdh23^+/−^). We found that the high-frequency response in both lines is restored and that the topographic map is fully restored for the F1(CBA/CaJ × C57BL/6) and partially restored for B6.cdh23^+/−^ mice. These results demonstrate that high-frequency hearing is required to form a topographic map of auditory space and that C57BL/6 mice have an early-onset hearing deficit which results in a loss of frontal RFs.

## Results

### C57BL/6 mice do not have a topographic map of auditory azimuthal space in the SC

To determine the spatial auditory receptive fields (RFs) of the SC neurons and their corresponding physical locations in the brain, we used virtual auditory space (VAS) stimuli and large-scale silicon probe electrophysiology to record from awake, head-fixed, 2-4 month old C57BL/6 mice (ordered from The Jackson Laboratory and bred in our facility) allowed to freely run on a cylindrical treadmill (see Methods). We presented direction-localized sound to the animal that ranged from −144° to 144° in azimuth (18° spacing), and 0° to 80° in elevation (20°spacing), totaling 85 sound locations in spherical space (where 0° is the front of the animal). The response properties of the SC neurons were measured with 256-channel multi-shank silicon probes, spike sorted, and processed to determine the RF.

We recorded the spiking activity of 2096 neurons from the dSC (300-1600 μm from the surface of the SC) from ten C57BL/6 mice. To compare these responses with those of CBA/CaJ mice, we also reanalyzed our previously published data from eight CBA/CaJ mice (Ito et al., 2020), in which 1615 dSC neurons were recorded.

In the C57BL/6 mice, (58 ± 2)% of all identified neurons significantly responded to the white noise auditory stimuli (see Methods), which is a slightly smaller percentage than that of the CBA/CaJ mice ((70 ± 3)%, p = 0.0032, ANOVA). To estimate the spatial RFs of these auditory responsive neurons, we fit the Kent distribution (Kent, 1982) to the directional auditory responses of these neurons and determined whether a neuron had a spatially localized RF by comparing the information explained by the Kent distribution and an alternative flat distribution (Bayesian information criterion (BIC), see Methods). In C57BL/6 mice, (11 ± 2)% of the auditory responsive neurons had spatially localized RFs, similar to CBA/CaJ mice ((12 ± 1)%, Figure 1C, p = 0.73, ANOVA).

**Figure 1.**
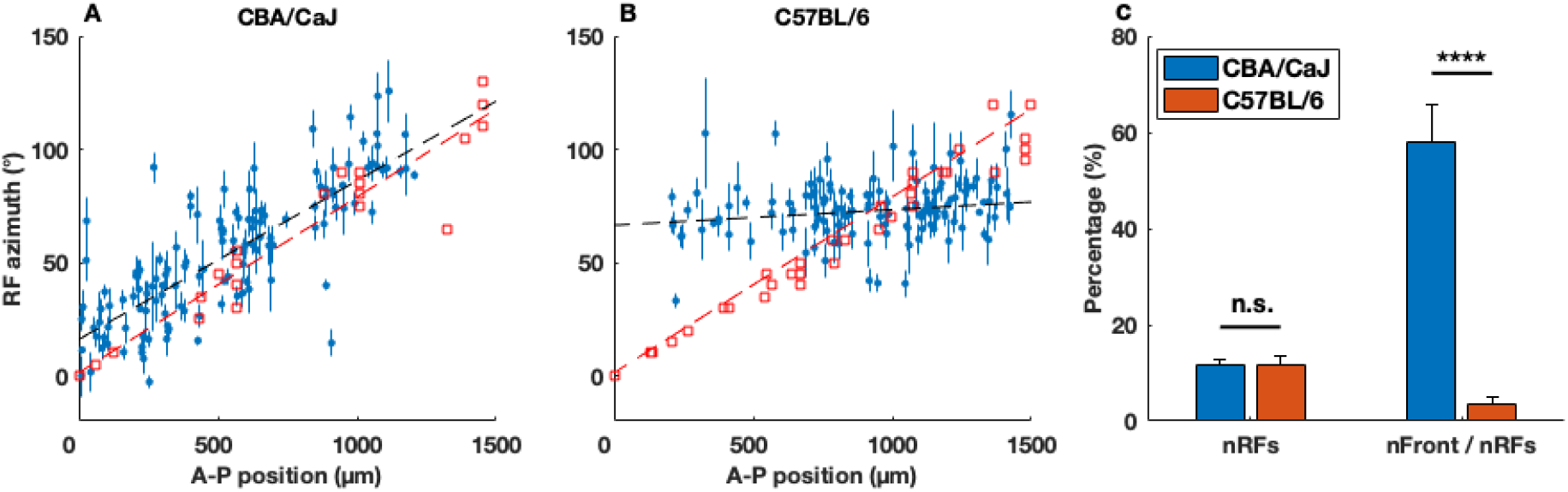
C57BL/6 mice lack a topographic map of the auditory azimuthal space along the anterior-posterior (A-P) axis of the superior colliculus (SC). ***A, B***. A plot of each SC neurons’ A–P positions vs. RF azimuths showing topographic organization along the A–P axis of the SC in CBA/CaJ (N = 8) but not in C57BL/6 (N = 10) mice. Each blue dot represents the auditory receptive field (RF) azimuth (center of the Kent distribution) and the A–P position of an individual neuron (error bars represent the statistical errors derived from the Kent distribution fits, systematic errors not included; see Methods); red squares are visual RFs measured by multi-unit activity in the superficial SC. The slope of the auditory RFs is (70 ± 6)° mm^−1^ for the CBA/CaJ mice and (7 ± 6)° mm^−1^ for the C57BL/6 mice (the black dashed line is the χ2 fit to the data, see Methods), and that of the visual RFs is (78 ± 4)° mm^−1^ for the CBA/CaJ mice and (75 ± 3)° mm^−1^ for the C57BL/6 mice (the red dashed line is the fit to the visual RFs, see Table 1 for comparison). The offset of the auditory RFs was (16 ± 4)° for the CBA/CaJ mice and (66 ±6)° mm^−1^ for the C57BL/6 mice. ***C***. There is no significant difference in the percentage of neurons with localized auditory RFs (BIC_Kent_ > BIC_flat_, see Methods) out of all auditory responsive neurons (nRFs, p = 0.73), but a significant difference in the percentage of neurons with a frontal RF out of all neurons with localized RFs (nFront / nRFs), in CBA/CaJ and C57BL/6 mice (****, p = 3 × 10^−16^). The error bars are SEMs of individual mice from each mouse strain.

The spatial auditory RFs of neurons in the dSC of CBA/CaJ mice are organized as a topographic map of space in which anterior dSC neurons respond to sound originating in the front of the animal and posterior neurons respond to sounds coming from the side. This auditory spatial map is aligned with the visual map in the superficial SC (Ito et al., 2021). To determine whether the SC in C57BL/6 mice as young as 2 months also contains an auditory map of azimuthal space that is aligned with the visual map, we calculated the correlation between the RF azimuth and the anteroposterior (A-P) position of the neurons in the SC. We found that, unlike in the CBA/CaJ mice (Figure 1A), most SC neurons with localized auditory RFs in C57BL/6 mice (Figure 1B) had a RF azimuth of ~70° (mean ± std: (74 ± 12)°), regardless of their A-P SC location (the A-P position vs RF azimuth plot for each CBA/CaJ and C57BL/6 mouse are shown in Figure 1-1). Using linear fitting to determine the slopes of the RF distributions vs. A-P location (see Methods), we calculated the slope of the auditory azimuthal map in C57BL/6 mice to be (7 ± 6) °/mm, which is significantly different from that of the visual map ((75 ± 3) °/mm, p = 4 × 10^−24^, calculated from multi-unit activity in the superficial SC), and that of the auditory map in CBA/CaJ mice((70 ± 6) °/mm, p = 5 × 10^−13^). This is because we find that C57BL/6 mice have far fewer neurons with frontal RFs (RFs with an azimuth < 57°, the mean and median azimuth of localized RFs for CBA/CaJ neurons in the exploratory data during the blind analysis, see Methods); there were (56 ± 6)% of neurons with frontal RFs in the CBA/CaJ mice, but only (4 ± 2)% in the C57BL/6 mice (p = 3 × 10^−16^, Figure 1C).

We note, as a cross-check, that in sharp contrast to the results for the auditory spatial maps, the visual spatial maps tell a different (but expected) story. The visual RF slopes and intercepts for the CBA/CaJ and C57BL/6 mice, as detailed in Table 1, are in complete statistical agreement.

**Table 1.** Quantitative comparison of receptive field properties and STRF features between mouse strains. N, number of mice of a strain; n, number of SC neurons; nRes, number of auditory responsive SC neurons; nRF, number of SC neurons with a localized RF; nFront, number of SC neurons with a frontal auditory RF; Slope/Intercept aud., the slope/intercept of the linear fit of A-P positions vs RF azimuths of SC auditory neurons; Slope/Intercept vis., the slope/intercept of the linear fit of A-P positions vs RF azimuths of the multiunit visual response in the superficial SC; nSTRF, number of neurons with a significant response to at least one frequency from 5 to 80 kHz in the random chord stimulus; >40kHz, number of neurons with a significant response to at least one frequency above 40 and up to 80 kHz in the random chord stimulus.

|  | CBA/CaJ | C57BL/6 | F1(CBA x C57) | B6.cdh23<sup>+/-</sup> |
| --- | --- | --- | --- | --- |
| N | 8 | 10 | 6 | 7 |
| n | 1615 | 2096 | 1266 | 1348 |
| nRes / n (%) | 70 ± 3 | 58 ± 2 | 74 ± 1 | 65 ± 4 |
| nRF / nRes (%) | 12 ± 1 | 11 ± 2 | 15 ± 3 | 12 ± 2 |
| nFront / nRF (%) | 56 ± 6 | 4 ± 2 | 51 ± 6 | 37 ± 5 |
| Slope aud.<br>(x 10<sup>-3</sup> °/mm) | 70 ± 6 | 7 ± 6 | 63 ± 5 | 38 ± 6 |
| Slope vis.<br>(x 10<sup>-3</sup> °/mm) | 78 ± 4 | 75 ± 3 | 79 ± 6 | 82 ± 5 |
| Intercept aud. (°) | 16 ± 4 | 66 ± 6 | 8 ± 4 | 31 ± 5 |
| Intercept vis. (°) | 1 ± 4 | -1 ± 3 | 0 ± 5 | -1 ± 4 |
| nSTRF / nRes (%) | 22 ± 1 | 27 ± 3 | 27 ± 3 | 32 ± 3 |
| >40kHz / nSTRF (%) | 98 ± 2 | 21 ± 9 | 99 ± 1 | 96 ± 2 |

### The SC of C57BL/6 mice does not respond to high-frequency sound

Previously, we showed that in the SC of CBA/CaJ mice, high frequency (40-80 kHz) spectral cues are used to form frontal RFs of azimuthal auditory space. Based on this, we hypothesized that the reason C57BL/6 mice lack frontal RFs is due to an inability of the mouse SC to detect high-frequency sound.

To determine the frequency tuning features of the auditory SC neurons in young C57BL/6 mice we measured the STRFs of the SC auditory responsive neurons by presenting the animal with uncorrelated dynamic random chord stimuli to each ear and calculated the corresponding spike triggered average (STA) of their responses (Ito et al., 2020). We are able to measure the frequency tuning of each neuron from the STRFs of individual SC neurons. In the C57BL/6 mice, we analyzed 323 neurons with significant STRFs, comprising (27 ± 3)% of the neurons that responded to sound stimuli, similar to that found in CBA/CaJ mice ((22 ± 1)%, p = 0.19). Of all the neurons with significant STRFs, only (21 ± 9)% had a significant response to sound frequencies above 40 kHz, which is drastically different from CBA/CaJ ((98 ± 2)%, p = 1 × 10^−17^, Figure 2E).

**Figure 2.**
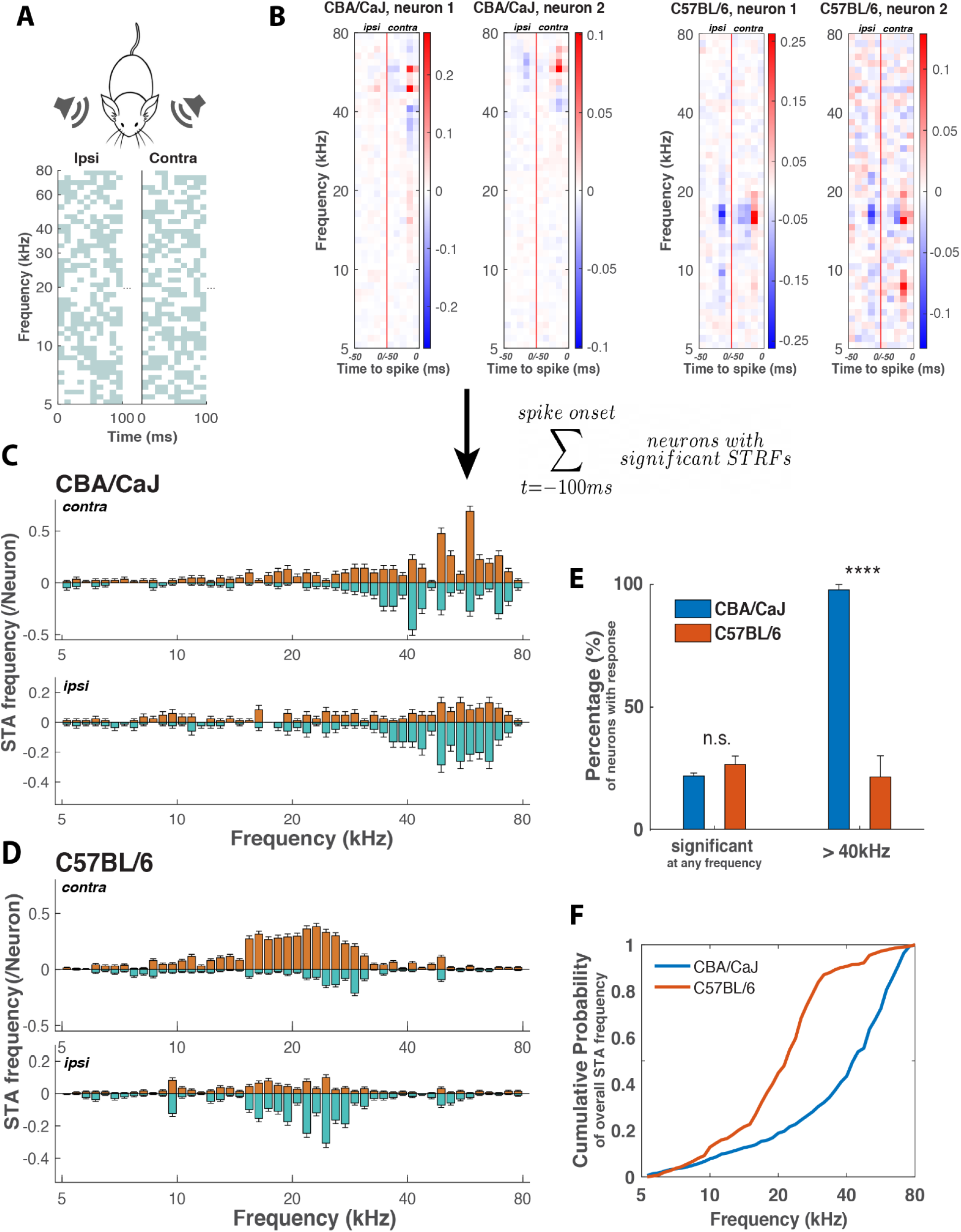
Auditory responsive neurons in the SC of C57BL/6 mice lack high-frequency responses. ***A***. A schematic representing the dynamic random chord stimuli. Top: independent random chord stimuli were played to each ear of the mouse. Bottom: an example of the first 100 ms of the dynamic random chord stimulus to the ipsilateral (ipsi; left) and contralateral (contra; right) ears. Grey pixels indicate the presence of a tone, with each time-frequency pixel randomly selected with a probability of 50%. The pixel time width is 10 ms, and the frequency ranges from 5-80 kHz, with 12 tones per octave. ***B***. Spectral temporal receptive fields (STRFs) of two example neurons from CBA/CaJ (left) and C57BL/6 (right) mice. For each example neuron, the spike-triggered average (STA) is calculated for the random chord stimuli played to the contralateral (right side of the red vertical line) and the ipsilateral ears (left side of the vertical line). The pixel color indicates the deviation from the average response at the corresponding frequency, showing the probability of a certain tone played on the contra/ipsilateral side to trigger a spike in a specific time bin for this particular neuron. ***C, D***. Pooled frequency tuning profiles from the STAs within 100ms prior to spike onset from all neurons with a significant STRF in CBA/CaJ and C57BL/6 mice (see Methods). For each strain, the top panel shows the contralateral frequency tuning profile and bottom the ipsilateral tuning profile. The orange color is the pooled positive response (significant red pixels from panel B) and the green color is the pooled negative response (significant blue pixels from B). ***E***. Fraction of neurons with significant response to any frequencies within the 5 to 80 kHz range and above 40kHz in the random chord stimuli in CBA/CaJ and C57BL/6 mice (****, p = 1 × 10^−17^). ***F***. Cumulative frequency tuning probability derived from C (CBA/CaJ) and D (C57BL/6).

To determine the collective frequency tuning features of all SC auditory neurons, we pooled the STAs of all neurons with a significant STRF, integrated over the 100 ms time window before the spike, and plotted the probability of SC neurons to respond significantly as a function of frequency across the 5 to 80 kHz range (see Methods). The resulting frequency tuning plot shows the overall preference in the response to a specific sound frequency for the population of neurons in the SC. The frequency tuning plot of C57BL/6 mice shows distinct features from that of CBA/CaJ mice (Figure 2C and D). Instead of a strong preference for ~60kHz in the CBA/CaJ mice, the C57BL/6 neurons did not have much response above 40 kHz, but preferably responded to a range of frequencies ~20 kHz. This difference in preferred frequency range is shown in the cumulative probability plots (Figure 2F), where the frequency preference for both contralateral and ipsilateral input is combined (see Methods). From the cumulative distribution, the C57BL/6 strain shows a sharp rise from ~20 to ~30 kHz, followed by a plateau-like smaller slope in the high-frequency range, indicating there is not much response above ~40 kHz.

Alternatively, the cumulative probability plot for CBA/CaJ mice depicted a much more pronounced response above 40 kHz. With a bootstrapping method, we found a statistically significant difference in the frequency tuning features between C57BL/6 and CBA/CaJ mice (p < 10^−6^, ɑ = 0. 001 for the two-sample Kolmogorov-Smirnov, see Methods and Fig 2-1). Together, these data show that the auditory responsive SC neurons in C57BL/6 mice lacked response to ~60 kHz sounds, which was previously found to be especially crucial for neurons to have frontal RFs. This finding supports our model that the SC auditory neurons use high-frequency spectral cues to compute frontal RFs.

### The high-frequency response of SC neurons is restored in F1(CBA/CaJ × C57BL/6) mice

Previous studies have shown that the first generation (F1) of mice from a cross of CBA/CaJ mice with C57BL/6 mice rescues the age-related hearing loss of C57BL/6 mice, as judged by measurements of the auditory brainstem response (ABR) (Bowen et al., 2020; Camlin et al., 2017). To determine if F1(CBA/CaJ × C57BL/6) mice rescue the lack of high-frequency SC responses, we recorded from six F1(C57BL/6 × CBA/CaJ) mice, as above, and collected the spiking activity of 1266 neurons from the dSC in response to random chord stimuli followed by STA analysis as in Figure 2. In this dataset, we find that a similar percentage of neurons that respond to sound stimuli had a significant STRF response, compared with CBA/CaJ and C57BL/6 mice (see Table 1 for detailed numbers). Of all the neurons with significant STRFs, almost all of them ((99 ± 1)%) had significant responses to sound frequency higher than 40 kHz, which is significantly more than C57BL/6 mice (p = 7 × 10^−19^) and similar to that of CBA/CaJ mice (p = 0.78, Figure 3C).

**Figure 3.**
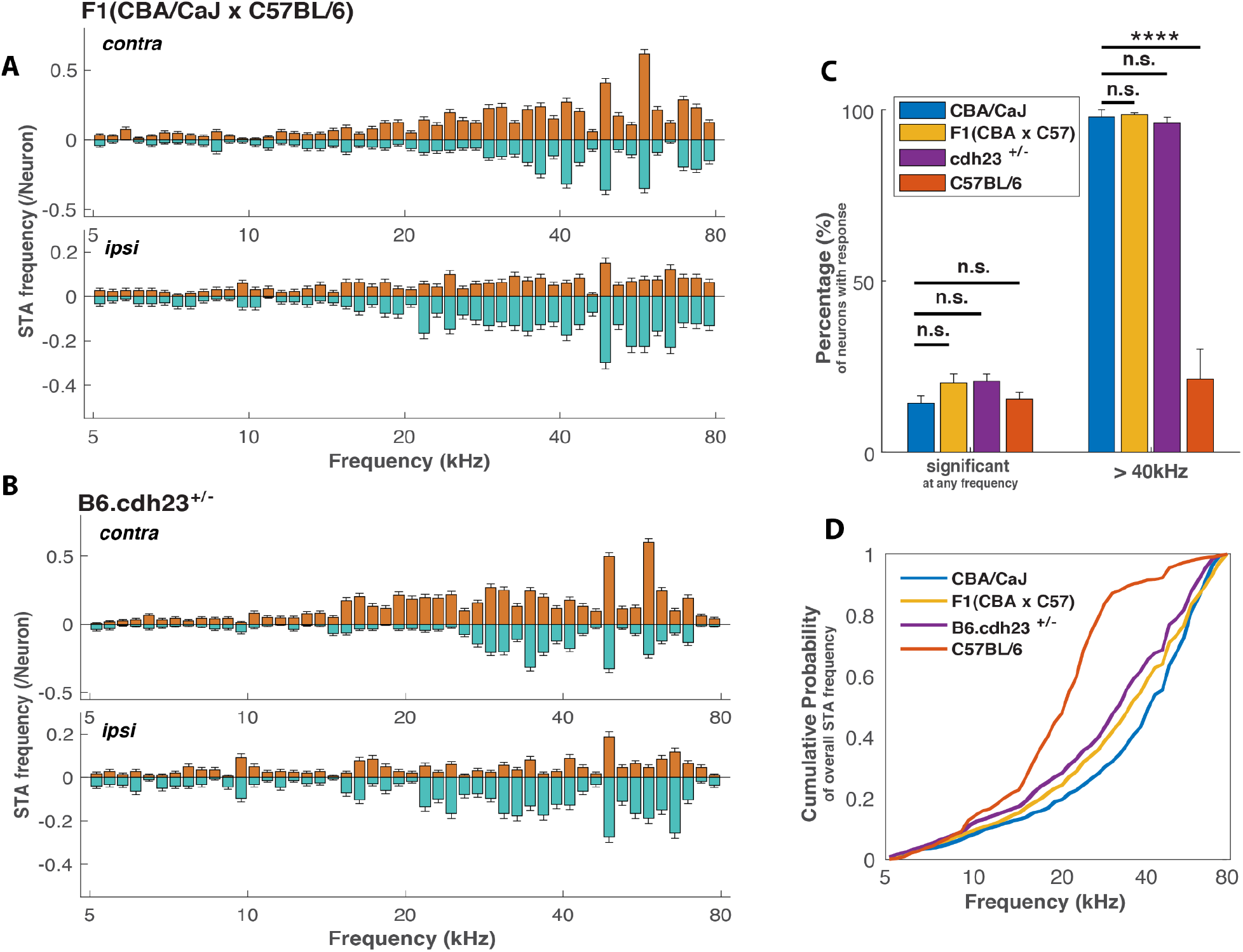
Frequency tuning profiles of F1(CBA/CaJ × C57BL/6) and B6.cdh23^+/−^ mice resemble that of CBA/CaJ but not C57BL/6 mice. ***A, B***. Pooled frequency tuning profiles from the STAs within 100ms prior to spike onset from all neurons with a significant STRF in F1(CBA/CaJ × C57BL/6) and B6.cdh23^+/−^ mice. Top, contralateral; bottom, ipsilateral. Orange color is the pooled positive response; green color is the pooled negative response. ***C***. Fraction of neurons with significant response to any frequencies from 5 to 80 kHz and above 40kHz in the random chord stimuli in F1(CBA/CaJ × C57BL/6 and B6.cdh23^+/−^ mice. (****, p = 3 × 10^−15^) ***D***. Cumulative frequency tuning probability derived from A (F1(CBA/CaJ × C57BL/6)) and B (B6.cdh23 ^+/−^) compared with CBA/CaJ and C57BL/6 mice.

Using the frequency plot to compare the frequency preference between strains, we found that the F1(CBA/CaJ × C57BL/6) mice respond to the random chord stimuli across all frequencies with an emphasis of sounds ~60 kHz, resembling what we found in the CBA/CaJ mice (Figure 3A). The cumulative distribution of frequency response reveals differences between the F1(CBA/CaJ × C57BL/6) mice and CBA/CaJ or C57BL/6 mice more directly: unlike the C57BL/6 mice, the F1(CBA/CaJ × C57BL/6) mice responded to > 40 kHz frequencies, but with a smaller emphasis than the CBA/CaJ mice. With the bootstrapping two-sample Kolmogorov-Smirnov test (see methods, Fig 4-1), we found that the frequency response distribution of F1(CBA/CaJ × C57BL/6) and C57BL/6 mice are statistically different (p < 10^−6^), but the same test failed to reject the null hypothesis that the distribution of the F1(CBA/CaJ × C57BL/6) and CBA/CaJ mice are indistinguishable (p = 0.94). This indicates that the SC auditory neurons of F1(C57BL/6 × CBA/CaJ) mice and CBA/CaJ mice have a similar frequency response in the SC; therefore, the CBA/CaJ genetic background is able to restore the high-frequency response in the C57BL/6 mice. This result is consistent with the hypothesis that C57BL/6 mice carry a recessive genetic mutation(s) that leads to a high-frequency response deficit in the SC.

**Figure 4.**
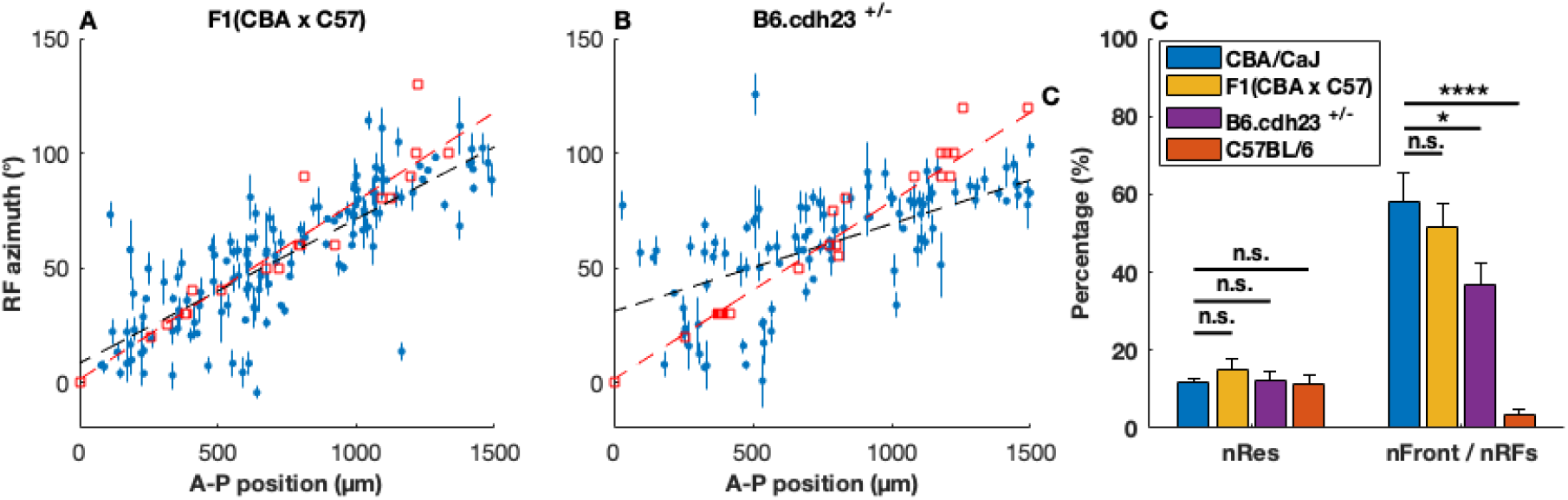
Topographic organization of RFs of SC auditory responsive neurons in F1(CBA/CaJ × C57BL/6) vs B6.cdh23^+/−^ mice. ***A, B***. A scatter plot of SC neurons’ A–P positions vs. RF azimuths showing topographic organization along the A–P axis of the SC in F1(CBA/CaJ × C57BL/6) (N = 6) and B6.cdh23^+/−^ (N = 7) mice (color and shape schemes same as Figure 1, see Table 1 for the slopes and intercept of visual/auditory linear fitting results). ***C***. Percentage of neurons with localized RFs (BIC_Kent_ > BIC_flat_, see Methods) out of all the auditory responsive neurons (nRFs), and percentage of neurons with a frontal RF out of all neurons with localized RFs (nFront / nRFs), in F1(CBA/CaJ × C57BL/6) (p = 0.50) and B6.cdh23^+/−^ mice (p = 0.024). The error bars are SEMs of individual mice from each mouse strain. (*, p = 0.02; ****, p = 3 × 10^−16^)

### The deficit of high-frequency SC responses of C57BL/6 mice is restored by the introduction of one wild-type *Cdh23* allele

Mutations in *Cdh23* are associated with age-related hearing loss in humans and mice (Bolz et al., 2001; Zardadi et al., 2020). C57BL/6 mice carry a point mutation in *Cdh23*, and the reintroduction of a wild-type *Cdh23* gene into C57BL/6 mice restores the age-related hearing loss, as determined by ABR experiments (Kenneth R. Johnson et al., 2017). To determine whether a single copy of wild-type *Cdh23* is sufficient to restore the high-frequency response in the SC of C57BL/6 mice, we recorded the responses of 1348 deep SC neurons from seven B6.cdh23+/− mice (Jackson labs) to random chord stimuli and measured their frequency tuning properties. We found that (32 ± 3)% of the auditory responsive neurons had significant STRF responses, similar to the other strains. Out of all the neurons with significant STRFs, most had significant responses to sound frequency higher than 40 kHz ((96 ± 2)%), which is significantly more than C57BL/6 (p = 3 × 10^−17^) but not statistically different from CBA/CaJ mice (p = 0.51).

The frequency plot shows that B6.chd23^+/−^ mice respond to random chord stimuli across all frequencies, resembling the response in CBA/CaJ and F1(CBA/CaJ × C57BL/6) mice. From the cumulative probability of frequency response we can see that unlike the C57BL/6 mice, the B6.chd23^+/−^ mice responded to > 40kHz frequencies, but with a smaller percentage than the CBA/CaJ or F1(CBA/CaJ × C57BL/6) mice. With the bootstrapping Kolmogorov-Smirnov method, we found that the frequency response distributions of B6.chd23^+/−^ mice and C57BL/6 mice are statistically different (p < 10^−6^), the same test failed to show that the distributions of the B6.cdh23^+/−^ and CBA/CaJ mice are significantly different from one another (p = 0.43, see methods and Fig 2-1). This suggests that the SC auditory neurons of B6.chd23^+/−^ mice have frequency responses more similar to the CBA/CaJ than the C57BL/6 mice, demonstrating that a single copy of the wild-type *Cdh23* gene is able to restore the high-frequency deficits of C57BL/6 mice.

### Restored frontal RFs and topographic organization in F1(CBA/CaJ × C57BL/6) and B6.cdh23^+/−^ mice

The rescue of the high-frequency response of C57BL/6 mice predicts that these mice will be able to process the information required to form frontal RFs and thus restore the topography of the auditory map of space in the SC. In order to test this, we also recorded from the F1(CBA/CaJ × C57BL/6) and B6.cdh23+/− mice and collected the spiking activity of neurons from the dSC while presenting the animal with spatially restricted white-noise stimuli.

In the F1(CBA/CaJ × C57BL/6) mice, we observed that (74 ± 1)% of all neurons were auditory responsive, out of which (15 ± 3)% of these neurons had spatially localized RFs, with (51 ± 6)% having a frontal RF. This is about the same as CBA/CaJ mice (p = 0.59) but significantly different from C57BL/6 mice (p = 1 × 10^−14^, Figure 4C, see Table 1 for detailed numbers). This indicates that the frontal RFs are indeed restored in the F1(CBA/CaJ × C57BL/6) mice. To determine whether the topographic auditory map is restored, we plotted the RF azimuth vs AP position for all the neurons with a localized RF and observed a positive correlation between the AP positions of these neurons and their auditory RF azimuths, as is shown in Figure 4A (The topographic maps for each F1(CBA/CaJ × C57BL/6) mouse are shown in Figure 4-1). The slope from the linear fit is (63 ± 5) × 10^−3^ °/mm and the intercept (8 ± 4)°, which are not different from those from the CBA/CaJ mice (p = 0.39 and 0.20, respectively), but are significantly different from C57BL/6 (p = 2 × 10^−12^ and 9 × 10^−15^, see Table 1). This result demonstrates that the rescue of the high-frequency hearing loss correlates with the rescue of the auditory spatial map in the SC.

In the B6.cdh23^+/−^ mice, we observed that (65 ± 4)% of the recorded neurons were auditory responsive, and (12 ± 2)% of them had spatially localized RFs. Out of these neurons, (37 ± 5)% of them had frontal RFs. This percentage is smaller than that of the CBA/CaJ mice (p = 0.02) but significantly larger than C57BL/6 mice (p = 2 × 10^−9^). To determine whether the topographic auditory map is restored in this strain, we again plotted the RF azimuth vs AP position for all the neurons with a localized RF and observed that there is a positive correlation of the AP position of these neurons and their auditory RF azimuth, as is shown in Fig 4B (the topographic maps for each B6.cdh23^+/−^ mouse are shown in Figure 4-1). The slope from the linear fit is (38 ± 6) × 10^−3^ °/mm and the intercept (31 ± 5)°, which is significantly different from those from the CBA/CaJ mice (p = 2 × 10^−4^ and 0.02) and C57BL/6 (p = 1 × 10^−4^ and 8 × 10^−6^). This suggests that one wild-type copy of *Cdh23* in the C57BL/6 genetic background partially rescues the topographic organization of auditory responsive neurons in the SC.

## Discussion

In this study we show that 2-month-old, C57BL/6 mice lack the ability to detect the high-frequency (>40kHz) spectral cues required for SC neurons to form frontal RFs; consequently, C57BL/6 mice do not have a topographic map of the auditory space in the SC. We also found that the high-frequency response deficit and the topography of the auditory map of space are both restored by either crossing the C57BL/6 with CBA/CaJ mice, or by introducing one copy of the wild-type *Cdh23* into the C57BL/6 genetic background. Taken together, these results show that high-frequency hearing is required to form a spatial map of auditory space along the azimuthal axis.

By recording from SC neurons in awake-behaving mice in response to VAS stimulation, we found that auditory neurons in the 2-4-month-old C57BL/6 mouse SC do not form an azimuthal topographic map of auditory space, with most neurons tuned to ~ 70° azimuth and a slope of azimuth across the A-P axis of the SC dramatically different from that of the visual map in the superficial SC or the auditory map of CBA/CaJ mice. Since C57BL/6 is one of the most widely used mouse strains in auditory research, this deficit could potentially be one of the reasons why the existence of an auditory topographic map of space in the SC of mice has only recently been shown. This finding should alert auditory researchers that C57BL/6 mice have significant high-frequency hearing deficits even at 2 months of age, before the loss of hearing sensitivity as measured by the auditory brainstem reflex (Li & Borg, 1991) or by low-frequency responses in A1 (Bowen et al., 2020).

Recessive mutations in *Cdh23* are associated with Usher syndrome type 1d in humans, whose symptoms include congenital hearing loss (Bolz et al., 2001; Zardadi et al., 2020). Cdh23 is a component of the tip-link in hair-cell stereocilia (Siemens et al., 2004), and studies have shown that the mutation of *Cdh23* leads to the deterioration of hair-cell stereocilia over time (Hequembourg & Liberman, 2001; Li & Borg, 1991). C57BL/6 mice are homozygous for a recessive mutation in *Cdh23*, which is postulated to account for much of the age-onset deafness observed in the strain (K. R. Johnson et al., 1997; Noben-Trauth et al., 2003). However, the early onset and the high-frequency aspects of this hearing deficit have not been reported except for a brief mention in 1980 (Henry & Chole, 1980). This may be because previous studies used ABR to evaluate the progression of hearing loss, but only used pure tone or click stimuli up to 32 kHz in their evaluation (Hequembourg & Liberman, 2001; Kane et al., 2012). In our experiments, we measured the STRFs of SC auditory neurons using random chord stimuli covering the frequency range of 5-80 kHz. This allowed us to determine the frequencies required to make an individual neuron respond, and determine the frequency tuning properties of the population of auditory neurons. We find that in CBA/CaJ mice, SC auditory neurons detect sounds across the 5-80 kHz range but very few neurons are tuned to high frequency (>40 kHz) in C57/BL6 mice. It is precisely these frequencies that we identified as being used to form frontal RFs in CBA/CaJ mice (Ito et al., 2020).

Because most mouse genetic tools such as knockouts and transgenic mice are created in a C57BL/6 background, there is great interest in being able to restore the hearing deficit of this strain. Previous work has shown that the early-onset hearing loss of C57BL/6 mice can be rescued in two different ways. One way is to perform a cross C57BL/6 mice (that, for example, carry a specific Cre-line) with CBA/CaJ mice and assay the F1 mice from this cross. It has been shown that this rescues the age-related hearing deficit of C57BL/6 mice (Firisnia et al 2001, (Lyngholm & Sakata, 2019), demonstrating that it is recessive alleles carried by the C57BL/6 mice that lead to deafness. The auditory deficits of C57BL/6 mice can also at least be partially restored by changing the C57BL/6 mutant *Cdh23* allele with the wildtype allele using CRISPR (Mianné et al., 2016), although one report suggested that hearing is only partially restored in this mouse line (Burghard et al., 2019).

Here we show that each of these methods also restores the high-frequency auditory response in the 2-4-month-old SC, the loss of neurons with frontal RFs, and the auditory map of azimuth along the A-P axis of the SC. This further demonstrates the correlation of high-frequency hearing and the creation of frontal RFs in the SC. F1 (CBA/CaJ × C57BL/6) mice have very similar auditory response properties and topographic organization of auditory RFs to the parental CBA/CaJ mice (Figure 1A). This result demonstrates that C57BL/6 mice carry a recessive mutation(s) that affects the ability of the SC to respond to high-frequency spectral cues needed to compute sound direction coming from the front. It also suggests that molecular genetic tools used to study auditory circuitry, such as cell type-specific Cre-lines, can be introduced into a mouse line with excellent hearing via a simple cross (Lyngholm & Sakata, 2019). We also show that the B6.cdh23^+/−^ mice have comparable high-frequency SC response to the CBA/CaJ mice, but only partially restored topographic maps. Others have also shown that the C57BL/6 mice heterozygous for the *Cdh23/Ahl1* mutation show age-related deficits in auditory temporal processing (Burghard et al., 2019). Furthermore, since both F1(CBA/CaJ × C57BL/6) and B6.cdh23^+/−^ are heterozygous for the *Cdh23* mutation, we speculate that the reason for the absence of a complete restoration might be that C57BL/6 mice harbor other mutations outside of *Cdh23* that affect high-frequency hearing, as has been reviewed (Bowl & Dawson, 2015).

Note that many studies that use transgenic mice perform their experiments on younger (2-4 month) C57BL/6 mice to get around the potential complications of age-related hearing loss in this strain. Here we show that this might not be a sensible solution if high-frequency components of a sound are potentially important in the study. Because we find that the organization of a spatial auditory map in the SC is abnormal in these mice, we predict that even at 2-4 months of age, C57BL/6 mice may have sound recognition or localization defects. A number of important naturalistic sounds for mice are in the high-frequency range, such as pup calls and mating vocalizations (Neunuebel et al., 2015; Ó Broin et al., 2018). This raises an interesting ethological question of whether and how C57BL/6 mice compensate for this high-frequency response deficit.

The relevance of high-frequency hearing loss to the formation/maintenance of spatial RFs in the SC has not been reported before. Here we demonstrate a correlation between the inability to detect high-frequency sound in the SC and the failure to form an auditory topographic map along the A-P axis of the SC. This finding is consistent with our model of how the RFs of auditory SC neurons are formed. Namely, neurons with frontal RFs are positively influenced by sound with a specific pattern of high-frequency (40-80 kHz) sound presented to the contralateral ear; neurons with lateral RFs are not as sensitive to this spectral pattern within a single ear but prefer to fire when a stimulus contrast is presented between the two ears (Ito et al., 2020). Therefore, mice that lack a high-frequency (40-80 kHz) response in SC neurons are unable to form frontal RFs, but can use the contrast between ears to form lateral RFs, and are therefore tuned to a spatial direction (~70° in azimuth) where the ILD is maximized.

We only analyzed the topographic mapping in the SC on the azimuthal axis because more is known about how the SC auditory neurons form their azimuthal RFs using ILD and spectral cues. It is still not validated whether the CBA/CaJ mouse SC contains a topographic map of the elevation on the medial-lateral axis (Ito et al., 2020) and whether such an elevation map, if it exists, requires different forms of spectral cues. We speculate that the elevation mapping could be disrupted if high-frequency hearing is involved in the elevation mapping, and future investigation is needed on this topic.

## Materials and Methods

### Ethics statement

All procedures were performed in accordance with the University of California, Santa Cruz (UCSC) Institutional Animal Care and Use Committee.

### Procedures

We used the mouse head-related transfer functions (HRTFs) to present virtual auditory space (VAS) stimuli and recorded the activity of the collicular neurons from head-fixed, alert mice using 256-electrode multishank silicon probes. Our experimental procedures were described previously (Shanks et al., 2016, Ito et al. 2017, Ito et al. 2020) and are detailed below.

### Auditory stimulation

#### HRTF measurement and VAS stimulation

The HRTFs of C57BL/6 (this study) and CBA/CaJ (Ito et al. 2020) mice were measured and the VAS stimuli were generated as previously described (Ito et al. 2020). The measured HRTFs for CBA/CaJ mice and the explanation for their use are available in the figshare website with the identifiers https://doi.org/10.6084/m9.figshare.11690577 and https://doi.org/10.6084/m9.figshare.11691615. The HRTFs for C57 mice were generated with the same methods. Briefly, in an anechoic chamber, a microphone (Bruel & Kjaer, 4138-L-006) was coupled with the back of the ear canal of the mouse (euthanized and decapitated) to record the response to a pair of Golay codes. To measure the HRTFs as a function of the incident angle, the coupled mouse head and microphone were rotated by a stage and tilted by a hinge that was coupled to a stepper motor. The HRTFs for 1010 grid points (10 elevations: 0–90° with 10° steps, and 101 azimuths: 0–180° with 1.8° steps) in the upper right quadrant of the mouse are measured. The head-related impulse response (HRIR) was reconstructed from the responses to the Golay codes. The earphone HRTF (eHRTF) was also measured and subtracted when generating VAS stimuli (see below) to avoid the additional frequency response due to the earphone setup.

VAS stimuli were generated as previously described (Ito et al., 2020). In brief, each stimulus sound was filtered by a zero-phase inverse filter of the ES1 speaker (Tucker-Davis Technologies), the eHRTF, and the measured HRIR. The VAS stimulus is then delivered pointing toward the animal’s ear canals. We tried to reproduce the physical configurations used for the HRTF measurements so that the acoustic effect induced by this setup is canceled by an inverse filter of the eHRTFs. Based on the similar shapes of the ears and head, we used the CBA/CaJ HRTF for experiments in CBA/CaJ and F1(CBA/CaJ × C57BL/6), and the C57BL/6 HRTF for experiments in C57BL/6 and B6.cdh23^+/−^.

#### Full-field white noise stimulus

The baseline stimulus pattern was 100-ms white noise with linear tapering windows in the first and last 5 ms. After applying the filters mentioned above, the full-field white noise stimulus contains grid points of five elevations (0-80° with 20° steps) and 17 azimuths (−144° to 144° with 18° steps), totalling 85 points in the two-directional field. The stimulus was presented every 2s and repeated 30 times per direction at the intensity of 50 dB SPL.

#### Random chord stimulus

To measure spectral tuning properties of the neurons with localized RFs, we used dynamic random chord stimuli and calculated the STA of the stimuli for each neuron. The stimuli consist of 48 tones ranging from 5 to 80 kHz (12 tones per octave). Each pattern was either 10 ms or 20 ms long with 3 ms linear tapering at the beginning and the end. In each pattern, the tones were randomly set to either ON or OFF with a probability of 0.5. The total number of tones per pattern was not fixed. The amplitude of each tone was fixed and set to be 50 dB SPL when averaged over time. One presentation of the stimuli was 2-min long and this presentation with the same set of patterns was repeated 20 times to produce a 40-min-long stimulus. Tones from the left and right speakers were not correlated with each other in order to measure the tuning to contrast between the ears. We did not use a specific HRTF for this experiment, but simply canceled the eHRTF so that the stimulus sound was filtered to have a flat frequency response near the eardrums.

### Mice

CBA/CaJ, C57BL/6 and B6.cdh23^+/−^ mice were either obtained from Jackson labs (stock# 000664 and 002756) or the offspring from the Jax mice, and the F1(CBA/CaJ × C57BL/6) are the first generation offspring from the CBA/CaJ and C57BL/6 cross. For each strain, we used 2-4-month mice of each sex.

### Electrophysiology

One day before the recording, a custom-made titanium head plate is implanted on the mouse’s skull to fix the mouse head to the surgery rig and recording rig without damaging or touching the ears. On the day of the recording, mice were anesthetized with isoflurane (3% induction, 1.2 - 1.5% maintenance; in 100% oxygen) and a craniotomy was made in the left hemisphere above the SC (approximately 1 mm by 2 mm in an elliptic shape, about 0.6 mm from the lambda suture). After recovering from anesthesia, the mouse was head-fixed onto the recording rig, where it was allowed to locomote on a rotating cylinder. A 256-channel, 4-shank silicon probe was inserted into the SC with the support of a thin layer of 2% low-melting-point agarose (in saline), and a layer of mineral oil was added on top to prevent the brain from drying. The silicon probe was aligned along the A-P axis. The multi-unit visual RFs were measured as the probe recorded from the superficial SC in all four shanks. After this measurement the probe was lowered until most of the channels were within the dSC (approximately just past the strong visual response in the superficial SC). Then we lowered and raised the probe by 100-120 μm to avoid the probe movement due to the effect of dragged tissue during recording. Recordings were started about 20-30 minutes after inserting the probe.

## EXPERIMENTAL DESIGN AND STATISTICAL ANALYSES

### Blind analysis

To validate our results and reduce potential false-positive findings, we performed blind analysis. First, we analyzed the spiking activity data from half of the neurons randomly sampled from each recording. After exploring the datasets and fixing the parameters in our analysis, we performed the same analysis on the other half of the data to test whether the conclusion holds true on the blinded data. All the results reported in this study passed a significance test in both the exploratory and the blinded data.

### Spike-sorting

We used custom-designed software for spike sorting, which was also used in our previous published work (Ito et al., 2017, 2020). Raw analog signals were high-pass filtered at ~313 Hz and thresholded for spike detection. Detected spikes were clustered based on the principal component analysis of the spike waveforms on the seed electrode and its surrounding electrodes. A mixture of Gaussians models was used for cluster identification.

### Estimating AP location of neurons

We estimated the relative position of the silicon probes using the visual RF positions measured in the superficial SC. When we inserted a silicon probe, we measured the positions of the visual RFs on each shank in the superficial SC using multi-unit activity. We extrapolated the visual RFs to find an A–P position where the visual RF azimuth was 0°, and defined this point as the zero of the A–P position. To estimate the A–P position of the auditory neurons we assumed the most superficial electrode of the silicon probe was at 300 μm in depth, which is justified by consistently stopping penetration at the position where visual responses predominantly disappear from all of the electrodes. We determined the insertion angle of the probe into the SC by measuring the angle between the electroprobe shanks and the visual response in the superficial layer of SC, which is justified by the post-recording observations of the insertion angle (Ito et al., 2020). The position of a neuron relative to the probe was determined by a two-dimensional Gaussian fit to its spike amplitude across multiple electrodes (Ito et al., 2017, 2020). We only analyzed neurons with a positive A–P position in order not to include neurons located outside the SC.

### Significance test for the auditory responses

We used quasi-Poisson statistics for significance tests of the auditory responses of individual neurons (Ver Hoef & Boveng, 2007), in order to deal with the overdispersion of the post-stimulus firing rate due to factors such as bursting of the neural activity, as described in our previously published work (Ito et al., 2020). The overdispersion parameter was estimated by the variance of the spike count divided by the mean, which should be 1 if the spiking activity of a neuron is Poissonian. To determine the significance of the response, we first estimated an overdispersion parameter and considered a response to be significant if the p value of the neuron’s spike count is below 0.001 (p = 1 − CDF(N), where CDF is a cumulative distribution function of the quasi-Poisson distribution and N is the spike count of the neuron).

### Function fit to estimate the azimuth of the RFs of the neurons

We used a maximum-likelihood fit of the Kent distribution (Kent, 1982) to estimate the azimuth of an RF. The Kent distribution is a spatially localized distribution in a two-dimensional directional space, and the equation of the Kent distribution is given by:

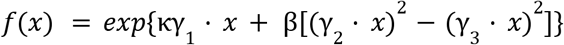

where × is a three-dimensional unit vector, κ is a concentration parameter that represents the size of the RF (κ > 0), β is an ellipticity parameter (0 < 2β < κ), the vector γ_1_ is the mean direction of the RF, and vectors γ_2_ and γ_3_ are the major and minor axes of the Kent distribution. These three unit vectors are orthogonal to each other. This function is normalized to 1 when × = γ_1_. At the limit of large κ, this distribution becomes asymptotically close to a two-dimensional Gaussian. At each point of the directional field, the likelihood value was calculated based on quasi-Poisson statistics (Ver Hoef & Boveng, 2007). The error of each parameter was estimated from the Hessian matrix of the likelihood function.

We used the front-Z coordinate system for the HRTF measurement and the stimulus presentation, but used the top-Z coordinate system for the Kent distribution fitting. We had to use the front-Z coordinate system for the HRTF measurement because of the restriction of our measurement stage. However, the front-Z coordinate system has a discontinuity of the azimuth across the midline that gives a problem in fitting and interpretation of the data near the midline. We avoided this issue by switching to the top-Z coordinate system. Because the Kent distribution in vector format is independent of the coordinate system, likelihood values and parameters in the vector format are not affected by this change when the fit is successful. To avoid an area where two coordinate systems are largely different, we only used neurons with their elevation smaller than 30° for azimuthal topography. To achieve stable fits, we restricted β to satisfy 4β < κ, set the azimuthal range to be from −144° to +144°, and set the elevation range to be from 0° to 90°.

### Estimation of additional systematic errors of the RF parameters

We estimated additional systematic errors that may affect the RF parameters as in (Ito et al., 2020). First, we did not consider the effect of locomotion on auditory neuron activities as a source of systematic error. Second, we take ±3° to be the estimated systematic error from the effect of eye movements. Third, we estimated the effect of the pinna movement on the HRTF as 13°. Fourth, our VAS stimuli might not be perfect because we used a single set of HRTFs for all the mice instead of measuring them individually. We quantified the overall difference (RMS) of HRTFs we measured from different mice and estimated the error as 5.3 ± 0.8 dB (Ito et al., 2020), and this corresponds to an angular shift of ±14°. To estimate the topographic map parameters, taking into account both the systematic and statistical errors, the systematic errors noted above were added in quadrature to give ±19°. In order to estimate the slope and offset parameters, we added, in quadrature, the estimated systematic error of ± 19° to each data point.

### Analysis of STRFs

We estimated the STRFs of neurons using the STA calculated from the spiking response to the dynamic random chord stimulus described above, as was done in our previous study (Ito et al., 2020). First, the spikes of each neuron are discretized in time bins the same as the stimulus segment size and the average of the stimulus patterns that preceded each spike was calculated. The stimulus was considered as 1 if there is a tone in the frequency and 0 otherwise. The significance tests for the STRFs were performed by examining whether the difference between the mean of the stimulus and the STA is significant. We used 0.001 as a significance threshold, and Bonferroni correction for evaluating multiple time bins, frequencies and contra/ipsilateral input. Neurons with <20 spikes during the stimulus were not analyzed.

When calculating the frequency tuning profiles of a certain mouse strain, all neurons with at least one significant STA pixel were considered as the total neurons with a significant STRF. At each frequency (5-80 kHz, 4 octaves, 12 tones per octave), all neurons with a significant STRF were evaluated whether there is a significant STA pixel within the 100ms time bin before spike onset. The frequency tuning plot shows, as a function of frequency, the probability of SC neurons with a significant STRF has a significant response at a particular frequency. And the error for each frequency is estimated assuming binomial distribution for each neuron to respond at a certain frequency.

We used a bootstrapping method and the Kirmogorov-Smirnov test to estimate the differences between mouse strains in terms of their collective cumulative frequency tuning curves. For each strain, all significant STA pixels (combining contralateral and ipsilateral, positive and negative, estimated as above) within 100 ms before spike onset were considered as a sample of frequencies that neurons from a mouse strain respond to. In each testing trial, 150 samples were randomly selected from each mouse strain, and a Kolmogorov-Smirnov test (alpha set to be 0.01) was done between all possible pairs of the strains. The number of trials such that the test successfully rejects the null hypothesis (that the samples from two strains are from the same distribution) is noted as an indicator (n_rej_) of how significant the differences of the frequency tuning curves between pairs of mouse strains are. A total of 1000000 trials were done between each pair (n_total_). The p value for each pair is then calculated as p = (n_total_ - n_rej_) / n_total_ × 100%. Since we only did the bootstrapping test for 1000000 trials, the p value is less than 10^−6^ if the null hypothesis was rejected in every trial tested. The testing results are illustrated in Figure 2-1.

## Acknowledgements

This work was supported by the National Institutes of Health (R01DC018580, R21 EY032230-01), and a donation from John Chen to A.M.L. We thank Soti ris Masmanidis for providing us with the silicon probes; and Jena Yamada and Brian Mullen for helpful comments on the manuscript.

Y.S., S.I., A.M.L. and D.A.F. conceived the project. Y.S. and S.I. designed the experiments. Y.S. acquired and analyzed the data. Y.S., S.I., A.M.L and D.A.F. interpreted the results and wrote the manuscript.

These authors jointly supervised this work: David A. Feldheim, Alan M. Litke

---

**Figure 1-1.**
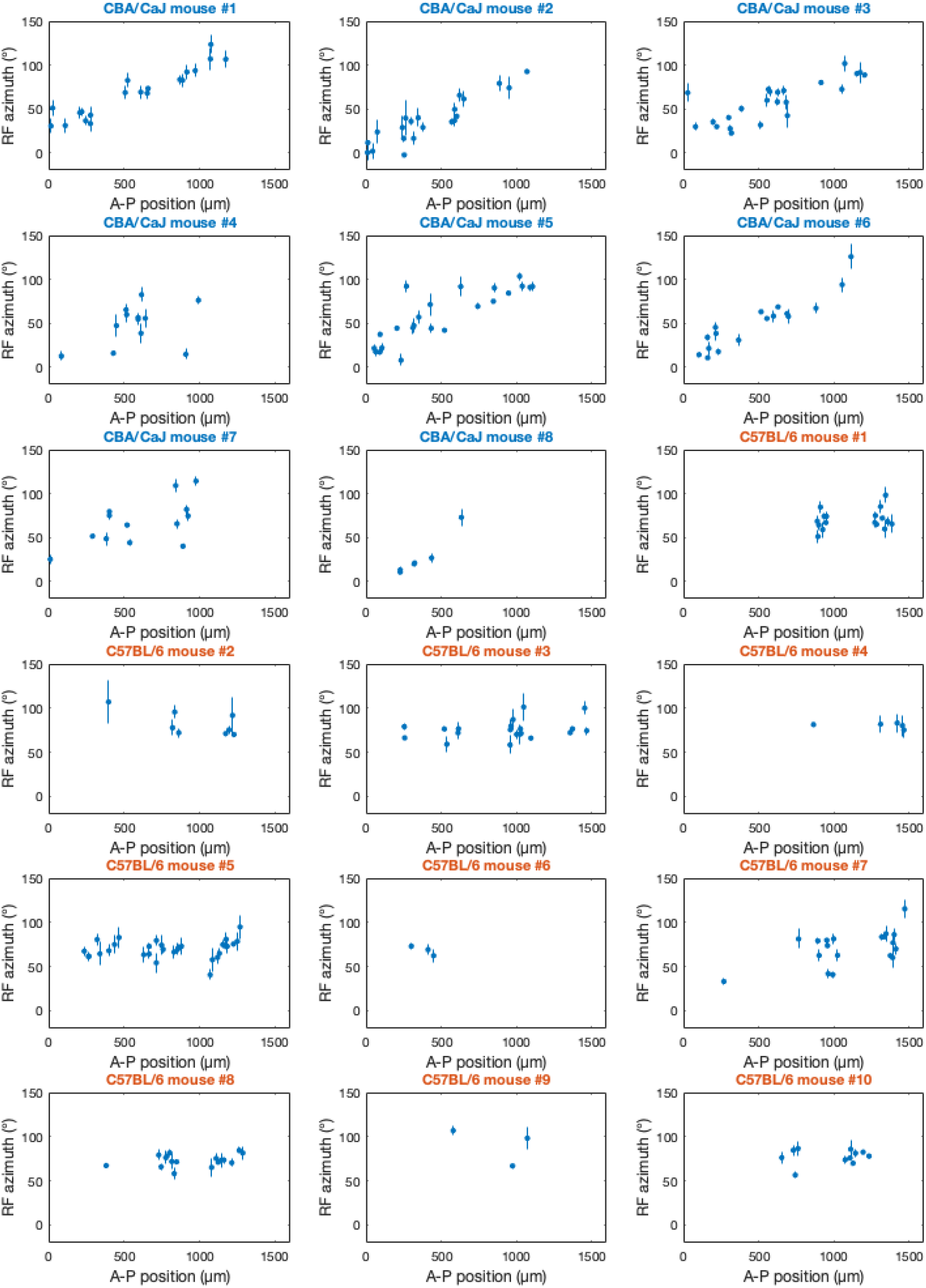
Topographic maps of auditory space in dSC from individual mice of the CBA/CaJ and C57BL/6 strain. Blue, CBA/CaJ mice; red, C57BL/6 mice.

**Figure 2-1.**
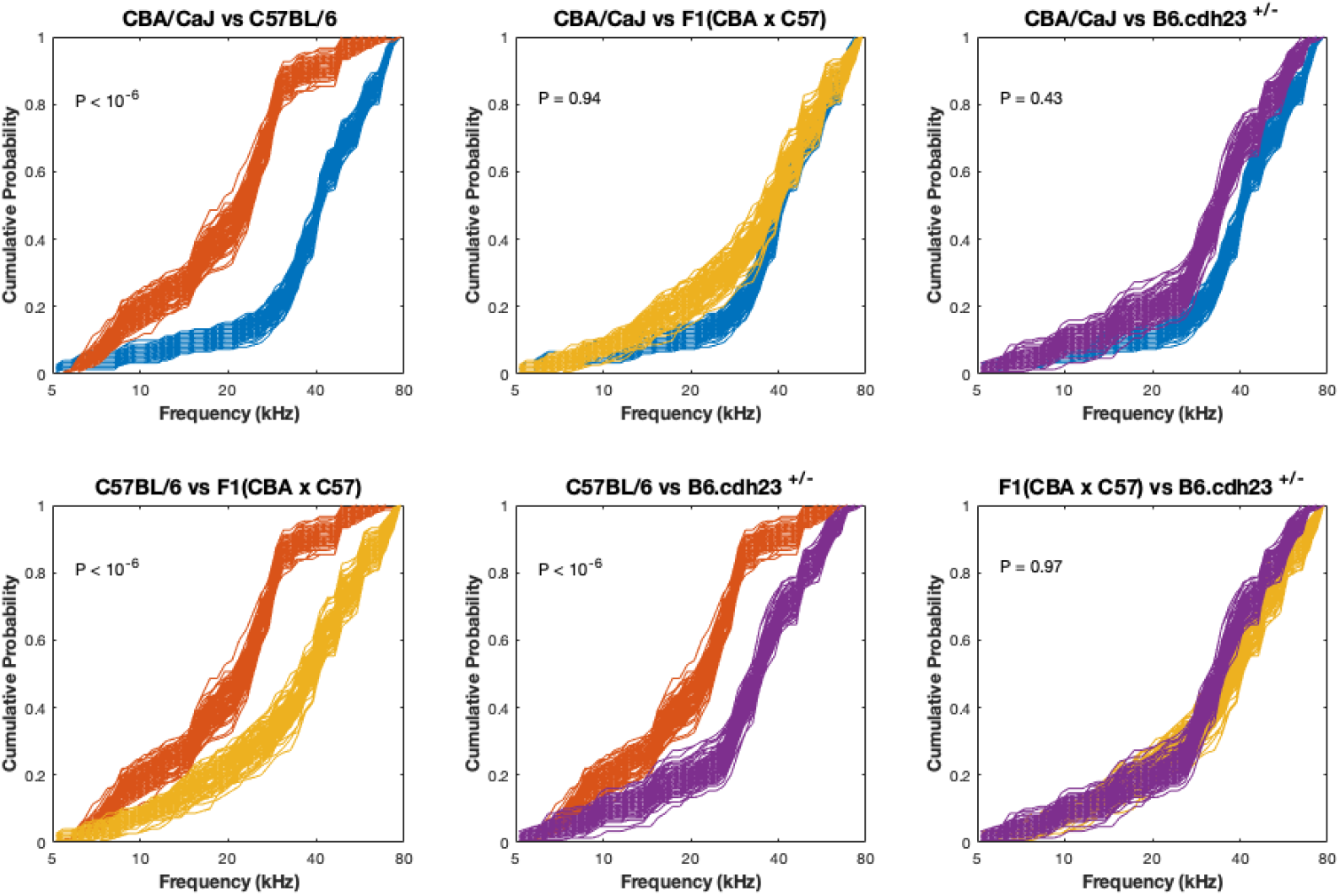
Results from bootstrapping Kolmogorov-Smirnov test for the cumulative frequency tuning curves. Each panel shows the comparison between a pair of strains. In each panel, 100 trials of the resampled data from the original data of each strain are plotted. Red, C57BL/6; blue, CBA/CaJ; yellow, F1(CBA/CaJ × C57BL/6); purple, B6.cdh23^+/−^.

**Figure 4-1.**
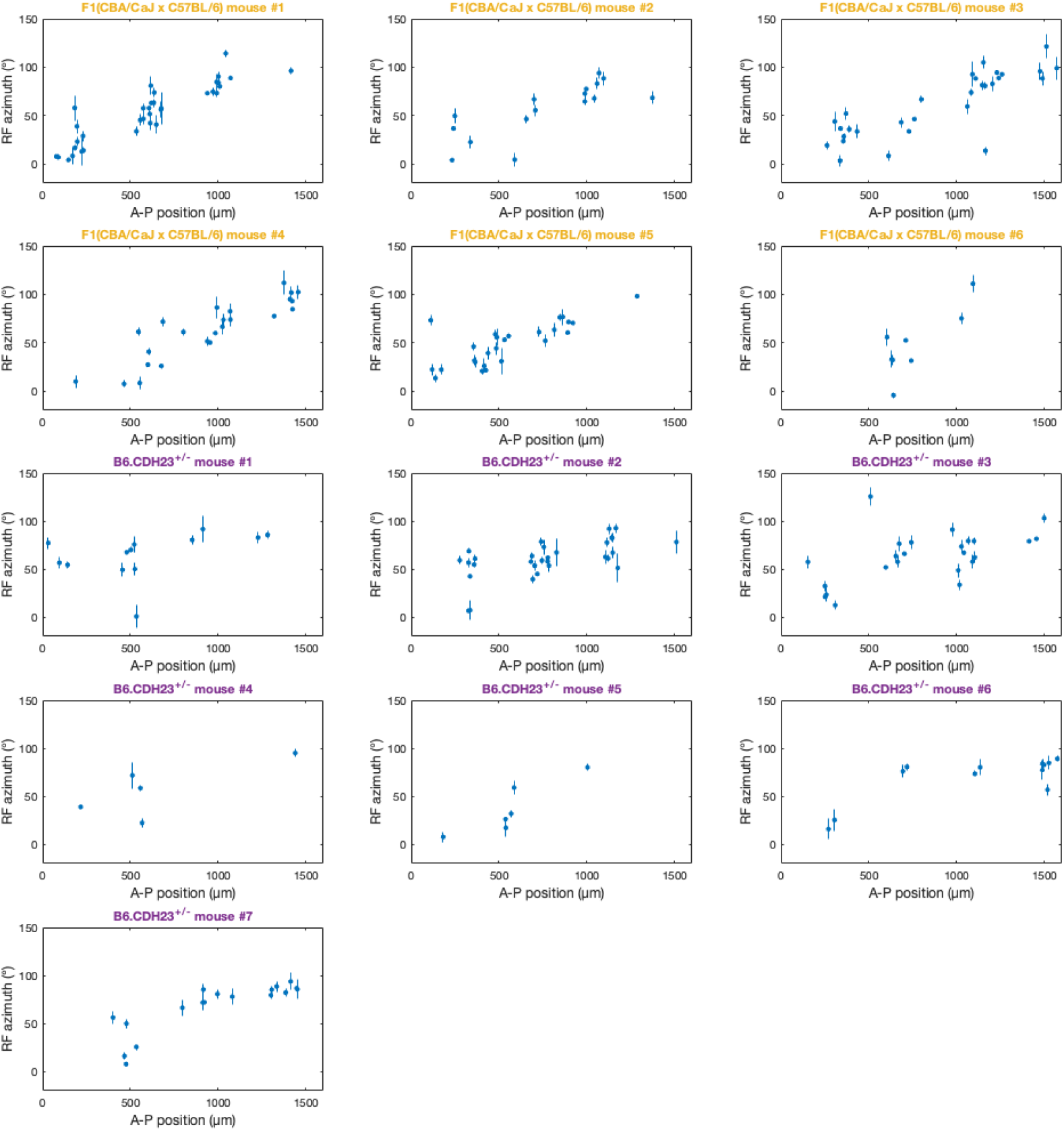
Topographic maps of auditory space in dSC from individual mice of the F1(CBA/CaJ × C57BL/6) and B6.cdh23^+/−^ strain. Blue, F1(CBA/CaJ × C57BL/6) mice; red,B6.cdh23^+/−^ mice.

## Notes

### Competing Interest Statement

The authors have declared no competing interest.

